# Defining the DNA Binding Specificity of GRHL2

**DOI:** 10.64898/2026.04.16.719077

**Authors:** Paige E. Messa, Christopher L. Warren, Noah R. Nicol, Keenan S. Pearson, Justin P. Peters, Amy M. Fowler, Elaine T. Alarid, Mary Szatkowski Ozers

## Abstract

Grainyhead-like 2 (GRHL2) is an epithelial transcription factor with context-dependent regulatory roles, yet the sequence rules governing its DNA recognition remain incompletely defined. In this study, a high-density genomic Specificity and Affinity for Protein (SNAP) DNA-binding array containing 772,732 tiled probes derived from GRHL2 ChIP-seq regions was used to resolve GRHL2 binding specificity at 6 base pair resolution across genomic sequences. From high-affinity probes, de novo motif analysis recovered the canonical 5’-AACCGGTT-3’ motif.

Sequence specificity landscapes revealed a stepwise reduction in binding as mismatches were introduced, with the strongest effects at the C (position 3) and G (position 6) within the motif, greater tolerance at the central CG dinucleotide, and intermediate tolerance at the A/T bases at the motif edges. This analysis also demonstrated the influence of nearby flanking sequences.

Extended motif and spacing analyses indicated dimeric binding at paired motifs, with periodic helical spacing consistent with interactions on the same face of the DNA helix. Integration of SNAP array binding with ChIP-seq data distinguished direct, motif-encoded GRHL2 occupancy from indirect, cofactor-mediated recruitment at genomic sites. These results define the sequence specificity of GRHL2 interactions with variations in the DNA consensus motif and flanking sequences within an endogenous genomic context.

**Graphical Abstract:** 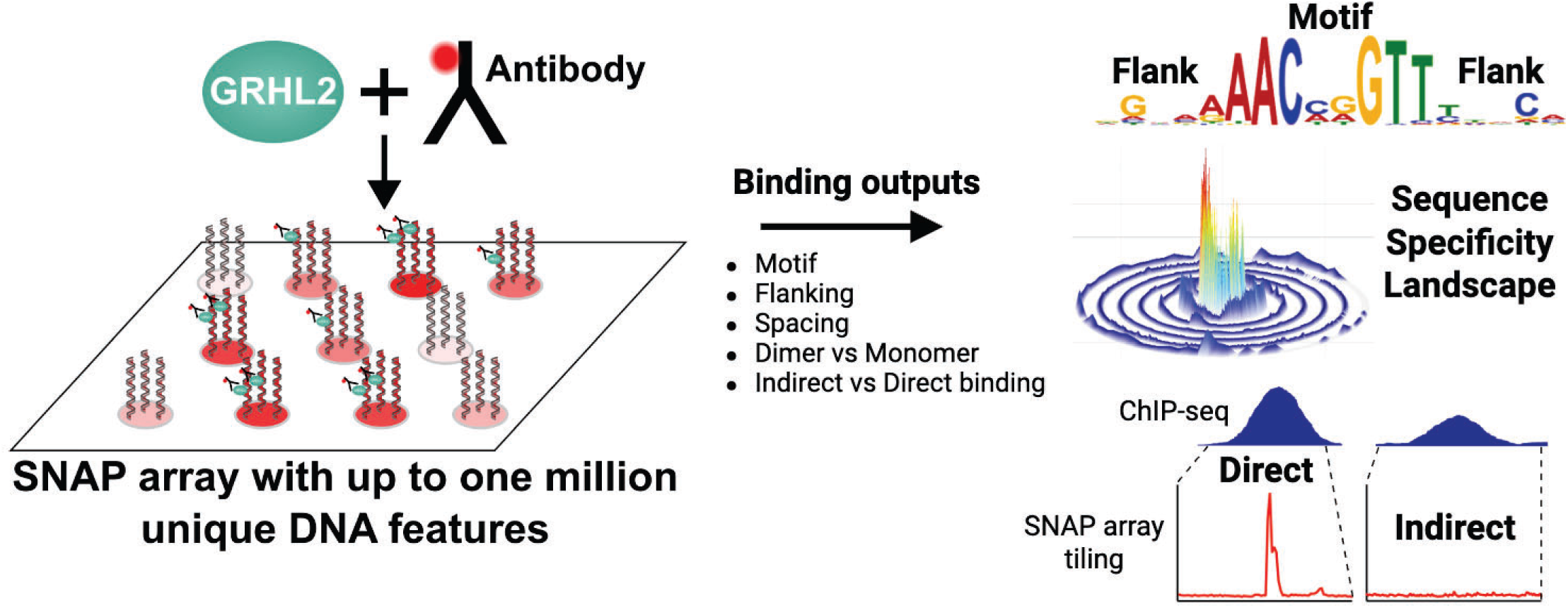

## INTRODUCTION

The Grainyhead-like (GRHL) protein family is a highly conserved family of nuclear transcription factors comprised of GRHL1, GRHL2, and GRHL3, each of which play vital roles in epithelial development and maintenance (1–4). Although GRHL family members share structural homology, including recognition of an identical DNA consensus sequence (5’-AACCGGTT-3’), individual family members provide distinct biological functions and possess spatial and temporal differences in expression during developmental stages (1). Loss of GRHL2 in mice causes severe defects in neural tube closure at anterior and posterior regions resulting in embryonic lethality (5, 6). Beyond its developmental roles, GRHL2 is a key regulator of epithelial cell identity that has been shown to exhibit a hybrid role in epithelial-to-mesenchymal transition (EMT), by inhibiting EMT but also by promoting cellular migration and plasticity in a process critical in cancer etiology (3, 4, 7–9).

GRHL2 can function as both a transcriptional activator and repressor depending on the cellular context (9–11). Genome-wide ChIP-seq studies across epithelial cell types have identified thousands of GRHL2 binding sites, and transcriptomic analyses following GRHL2 perturbation demonstrate regulation of hundreds of genes in a context-dependent manner (12–16). Many transcriptional effects appear to be indirect, meaning that they are mediated through a protein complex without direct contacts between GRHL2 and genomic DNA (17, 18). One such interacting partner has been identified as estrogen receptor a (ER) (19) which is expressed in over 70% of all breast cancers and is a major therapeutic target (20–23).

As the most abundant ERa post-translational modification, phosphorylation at S118 is associated with enhanced chromatin binding and estrogen-responsive gene expression, and its presence has been linked to endocrine therapy outcomes in ERa-positive tumors (24–28). GRHL2 motifs enriched at phosphorylated ERa (pS118-ERa) binding sites have been identified, suggesting a cooperating role between GRHL2 and pS118-ERa in regulating gene expression (29). Subsequent transcriptomic analysis revealed several genes co-regulated by ERa and GRHL2, as evidence of the ability of GRHL2 to both enhance and suppress ERa transcriptional activity (30). DNA binding motif analysis demonstrated significant enrichment of the GRHL2 motif in the tumor-specific ERa cistrome, and GRHL2 is essential for estrogen-stimulated proliferation in breast cancer cells (15). There is substantial evidence of tumor suppressive action by GRHL2 in a tissue-specific manner by prevention of EMT as well, but GRHL2 overexpression is paradoxically correlated with poor prognosis clinically (10). ERa and GRHL2 have also been shown to interact with FoxA1, which is of interest since both FoxA1 and GRHL2 have been shown to exhibit pioneer factor activity (19, 31, 32).

Before GRHL2 interactions with these other transcription factors can be deconvoluted, it is useful to first understand the biophysical binding characteristics of GRHL2 alone. Current ChIP-seq and epigenomic approaches are limited in their ability to resolve detailed binding determinants, flanking-sequence effects, and inter-site dependencies within GRHL2-bound regions. Because ChIP-seq measures protein occupancy in the presence of cofactors and chromatin structure, it cannot distinguish direct sequence-driven binding from cofactor-mediated interactions or parse the contribution of local sequence variation on binding specificity and affinity. Defining the GRHL2 binding preferences in detail will allow for the design of small molecules that directly target GRHL2-DNA interactions at precise sites to modify the expression of oncogenic factors.

In this study, GRHL2-DNA interactions are examined using genomic binding sites, as well as all permutations of monomeric and dimeric-spaced sites. Binding events were assessed using Specificity and Affinity for Protein (SNAP)-binding DNA microarray technology (29, 33, 34). The SNAP array is a custom genomic DNA array of up to 1 million features that is bioinformatically designed using overlapping peaks from four GRHL2 ChIP-seq data sets (12, 15, 30, 35), ultimately displaying 48-mer probes that tile across 6,151 regions at 6-bp spacing using 772,732 features of the array in the current configuration. This technology enables examination of GRHL2 binding across a wide range of genomic and control sequences in greater detail. Protein-DNA interactions are studied without the influence of other factors such as coactivators or co-binding proteins that may convolute the results. By integrating SNAP array binding measurements with ChIP-seq occupancy, direct, motif-directed GRHL2 binding can be differentiated from indirect recruitment to DNA to resolve the sequence characteristics that influence GRHL2 binding. In this study, we define specificity rules of GRHL2 motif recognition and examine how local sequence environment and motif organization contribute to GRHL2 binding to genomic DNA.

## MATERIALS AND METHODS

### GRHL2 SNAP Array Design

The SNAP array (Proteovista LLC), a custom genomic DNA array synthesized by Agilent Technologies, was bioinformatically designed to tile across GRHL2 binding sites based on overlapping peaks from four GRHL2 ChIP-seq data sets (12, 15, 30, 35). Overlapping peaks were merged into contiguous regions, and genomic intervals exceeding 1 kb were excluded. Of the 8,931 combined genomic regions identified that overlapped with all four ChIP-seq datasets, 6,151 were utilized to design DNA array probes tiling across at 6-bp spacing. Of these, 3401 regions were prioritized to include a subset with known GRHL2 binding motifs consisting of the consensus 5’-AACCGGTT-3’ site with up to one mismatch and regions shown previously by SNAP data (unpublished data) to have ERa/FoxA1 co-binding sites. The remaining 2750 regions were selected based on size. The probes consisted of 48 nucleotides from the genomic regions with a 12-nucleotide constant region on the 3’ surface-attached terminus used for primer extension. Control groups, comprised of 195,282 probes, included all permutations of the consensus site as a monomer and dimer, spacing variations between dimer sites, sites in forward and reverse orientations, and binding sites in conjunction with FoxA1 and ERa. A subset of 53,328 probes was designed as a single oligonucleotide that can form a hairpin of 25 bp (29, 33, 34) with a three-nucleotide loop, while the remaining probes were designed to be primer-extended to a length of 48 bp.

### Array Extension

To prepare the SNAP probes, the array was incubated in a conical tube of preheated 7 M urea in 1x phosphate-buffered saline (PBS) at 65 °C for 30 minutes, with brief shaking at 10-minute intervals. The array was then transferred to preheated 1x PBS and incubated at 65 °C for 15 minutes, with brief shaking at 7.5 minutes. Next, the array was incubated in non-stringent wash buffer (NSWB; 6x Saline-Sodium Phosphate-EDTA (SSPE) + 0.01% Tween-20 in ultra-pure water) for 15 minutes at room temperature covered from light before it was rinsed briefly in ice cold 1x Final Wash Buffer (FWB; 0.6x Saline Sodium Citrate (SSC)) and dried. Phusion DNA polymerase (New England Biolabs, M0530L) was used with a 12-nucleotide primer complementary to the constant probe region to extend the single-stranded genomic DNA probes using published procedures (36, 37), forming 60-bp double-stranded DNA probes. Proper extension was assessed on a control array using fluorescent Cy3-labeled dUTP (Cy3-dUTP; Supplementary Figure 1).

### Protein Binding and Detection

After primer extension, a hybridization chamber (Grace BioLabs, 623507) was mounted over the array, hydrated twice with ultrapure water, and blocked with 2.5% non-fat dry milk for 1 hour with rotation. Following blocking, arrays were rinsed with GRHL2 binding buffer (Proteovista LLC), and then 25 nM purified GRHL2 protein (Origene, TP314498) diluted in binding buffer containing 0.2% BSA plus Alexa Fluor 647-labeled anti-DYKDDDDK (e.g. FLAG) antibody (Cell Signaling, 15009S) was applied for 1 hour with rotation. The protein-antibody solution was then removed, and the array chamber rinsed with binding buffer. The array chamber was then removed, and the entire array briefly rinsed in a conical tube of ice cold 1x FWB, dried, and scanned using a GenePix 4000B scanner (Molecular Devices) and GenePix Pro 7 image acquisition software (Molecular Devices).

### Array Data Normalization

Raw Agilent microarray images were extracted with Agilent Feature Extraction Software (version 12.1). GRHL2 SNAP microarray experiments were performed in triplicate and normalized. Raw extracted data were globally normalized to account for overall differences in absolute data ranges between replicates and then quantile normalized to account for differences in intensity distributions (29). Finally, the median intensity of the three quantile normalized values for each probe was taken as its final normalized intensity. These final normalized intensities were used in all subsequent analyses.

## Data Analysis

### Dimer binding model

For features containing two GRHL2 binding motifs separated by defined numbers of base pairs, fluorescence as a function of motif spacing was fit with a DNA torsional elasticity-based helical phasing model (38–41). In this model, the periodic dependence of fluorescence on motif spacing is governed by the DNA helical repeat (h_r_) and is broadened by thermal torsional fluctuations, where s_min_ sets the phase offset of the first minimum. The predicted fluorescence at spacing s is:

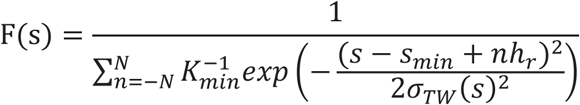

The summation was evaluated from n =-N to N, with N = 96, where h_r_ is the helical repeat that is defined as bp per turn, K_min_ is a scaling parameter corresponding to the effective binding strength at a phasing minimum, and a_TW_ is the standard deviation of the relative twist angle between motifs that increases with spacing due to DNA torsional elasticity.

Thermal broadening was modeled using:

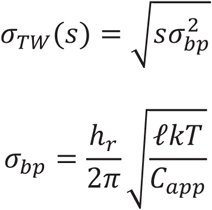

where C*_app_* is the apparent torsional modulus, f is the rise per base pair, k is Boltzmann’s constant, and T is temperature. Model parameters (h*_r_*, C*_app_*, K*_min_*, s*_min_*) were fit with nonlinear least-squares minimization to the forward-forward data using the nlminb function in R software (v4.4.3).

### Nucleotide/Motif Discrimination Analysis

For each position in the left flank (-5 to-1) and right flank (+1 to +5) of the consensus GRHL2 binding site, the base frequencies for the highest intensity quartile and lowest intensity quartile were calculated. The nucleotide enrichment ratios at each flanking position, as defined as the proportion of highest to lowest quartile frequencies, were then determined. Statistical significance across the four nucleotides at each position (p-value < 0.05) was determined with the chi-squared test (p-value < 0.05) at each flanking nucleotide position, wherein the enrichment ratio of each nucleotide at each position was compared to its overall background enrichment ratio across all genomic regions displayed on the SNAP array. For this analysis, only genomic probes with a perfect GRHL2 consensus site (5’-AACCGGTT-3’) were used. In addition, probes with a second perfect motif or one mismatch GRHL2 motif were excluded to avoid dimeric binding site effects. Finally, probes where the GRHL2 consensus site was within 10 bp of the primer site or end of the probe were excluded to avoid the effects of constant sequences or duplex terminus instability, respectively.

### Sequence Specificity Landscape (SSL)

SSLs were prepared as described (34, 42). For the flattened 2D SSL, a Python script was written to extract the sequences, x- and y-coordinates, and the intensities of the one mismatch ring from the tab-delimited text file underlying the 3D SSL. This data was then visualized in a two-dimensional format with lines separating the transitions between mismatch positions.

### Differential Binding of ERa and FoxA1 to SNAP Regions

The 6,151 SNAP regions were divided into two types: direct and indirect. This was determined by the probe with the maximal normalized SNAP intensity in each region. If the normalized intensity was greater than 3.0, then it was considered a GRHL2 direct binding region; otherwise, it was classified as an indirect GRHL2 binding region. Differential binding between the direct and indirect classes of SNAP regions using orthogonal methods were analyzed independently for ERa and FoxA1. The first differential analysis was based on the number of unique regions in each class that overlapped with a ChIP-seq peak. The second differential analysis was based on whether each SNAP region contained at least one genomic sequence for the transcription factor. The generalized half site 5’-RGGTCA-3’ (R = A or G) was used for ERa, and the consensus sequence 5’-AWTRTTKRYT-3’ (W = A or T; K = G or T; V = C or T) was used for FoxA1. The chi-squared test for independence with a p-value < 0.05 was used for each protein and each method.

## RESULTS

### Design and validation of a high-resolution genomic SNAP array for GRHL2 DNA binding sites

Towards the subsequent interrogation of GRHL2-DNA binding specificity, the domain organization of mammalian GRHL2 (isoform 1; NP_079191.2) was examined (Figure 1). GRHL2 is a transcription factor of 625 amino acids (aa) composed of an N-terminal transactivation domain (aa 1–133) and a central DNA-binding domain (DBD; aa 247–484) that belongs to the Grainyhead/CCAAT Box-Binding Protein 2 (CP2) family of winged-helix transcription factors (10, 43). Within the DBD, a conserved core segment (aa 416–424) corresponds to the helix-turn-helix element that forms some base-specific contacts with the GRHL consensus motif (5’-AACCGGTT-3’), as reported in prior structural studies (43). At the C-terminus, GRHL2 contains a dimerization domain (aa 494–625) that mediates stable homodimer formation.

**Figure 1.**
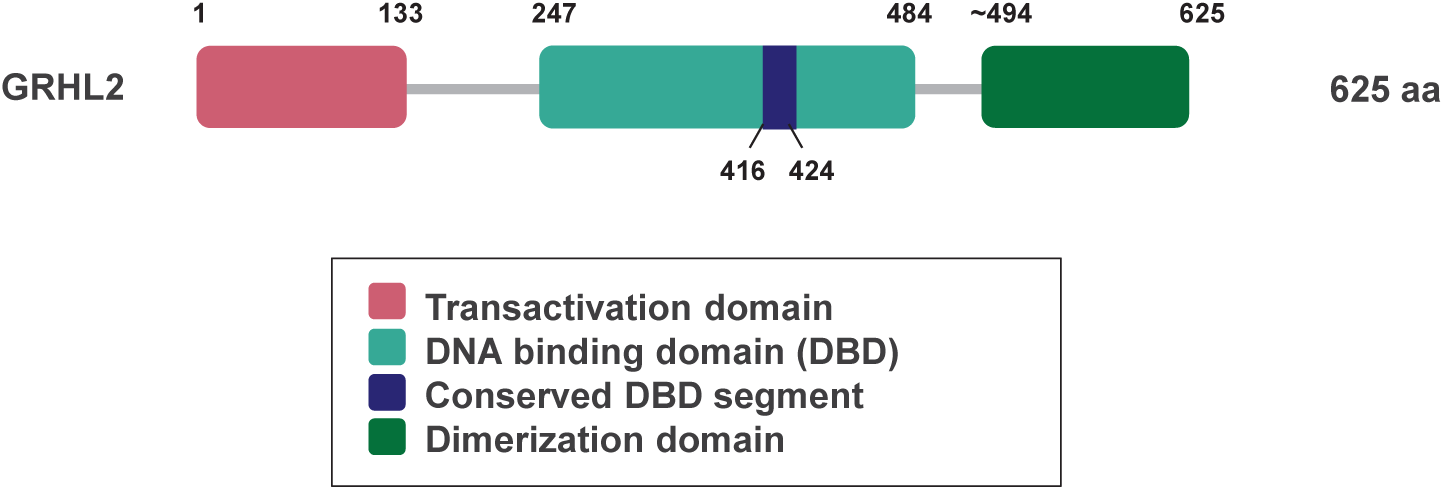
Functional domain organization of human GRHL2. Domain structure of GRHL2 (isoform 1) (Accession number NP_079191.2) showing the transactivation domain, DNA-binding domain (DBD), conserved DBD segment, and dimerization domain, as adapted from (10).

To systematically define the sequence determinants of GRHL2-DNA recognition across genomic sequences, we generated a custom high-density SNAP DNA-binding microarray composed of a total of 968,014 DNA features with 772,732 unique double-stranded DNA features derived from GRHL2 ChIP-seq datasets (Figure 2A). Conserved genomic target regions were selected from four independent GRHL2 ChIP-seq datasets (12, 15, 30, 35). Although each dataset possessed thousands of GRHL2-associated peaks, only a subset of peaks (8,931) was shared across all four experiments due to context-specific (e.g. cell type and experimental conditions) occupancy (Figure 2B). To capture shared binding regions, overlapping peaks were merged, and the resulting genomic intervals were tiled at 6-bp spacing to create sets of related probe sequences that differ only by local sequence shifts. The remaining 195,282 DNA features of the array were occupied with control probes and other altered sequences designed to test binding specificity, including probes where GRHL2 consensus and mismatch sites were combined at various spacings with ERa and FoxA1 consensus and mismatch sites. Since ChIP-seq peaks span hundreds of base pairs and GRHL2 does not bind evenly throughout the peak, the 6-bp tiling design enabled fine dissection of flanking sequence contributions, inter-motif spacing, and local binding preferences within individual ChIP-seq loci. Purified GRHL2 protein was added to the array in the absence of cellular cofactors, and bound features were detected using an anti-FLAG fluorescent antibody for quantitative measurement of both sequence specificity and affinity across the tiled genomic sequences (Figure 2C).

**Figure 2.**
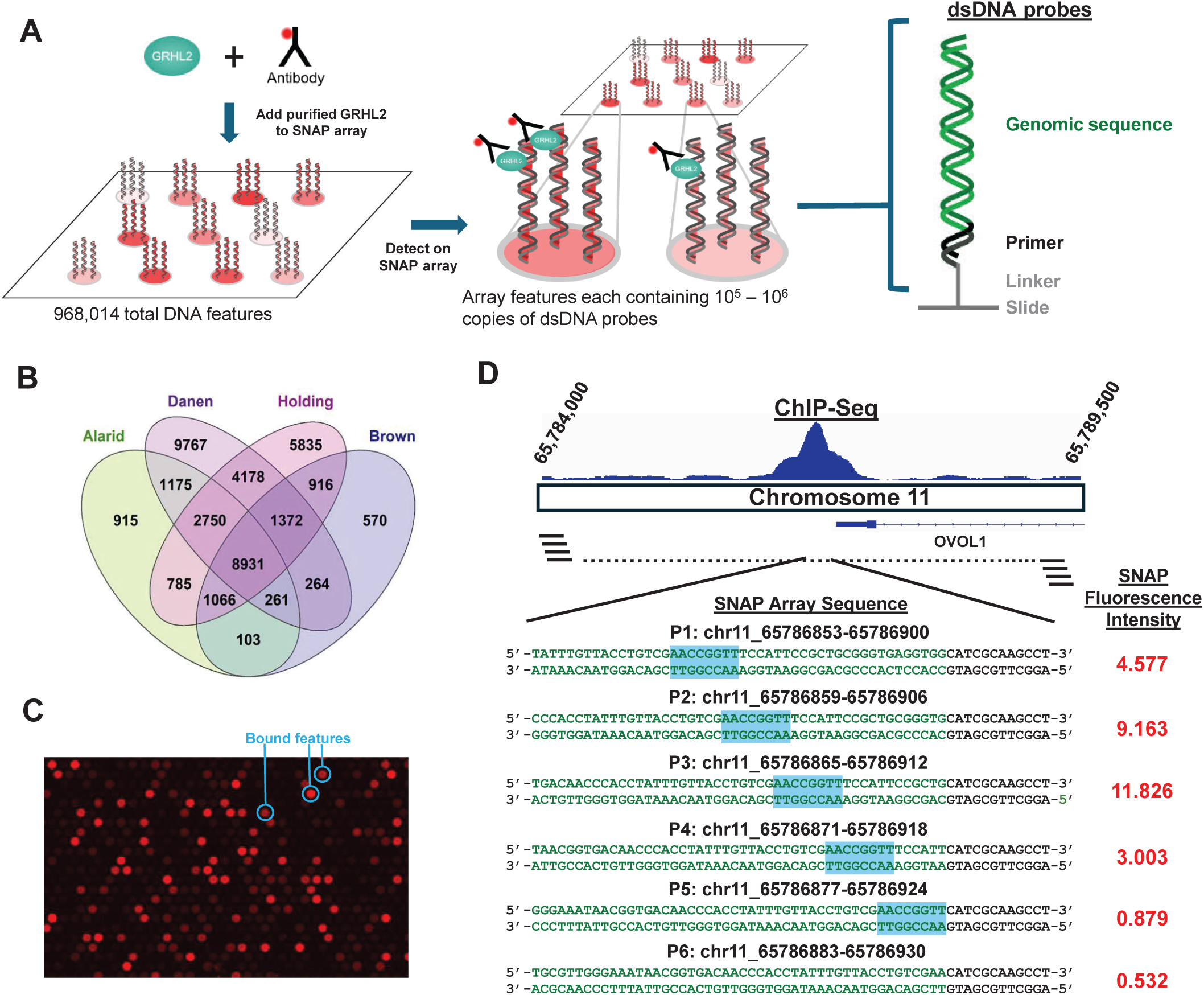
Design of the SNAP array enables high-resolution measurement of GRHL2 binding across genomic ChIP-seq regions. **A)** Left: Purified GRHL2 protein and fluorescently-labeled GRHL2 antibody assayed on the SNAP array. Middle: Subsection of the GRHL2 array displaying GRHL2 with antibody bound to DNA polymerase extended double-stranded probes. Right: Probes consisted of variable 48-nt genomic regions (green) with a 12-nt constant region (black) on the 3’ end used for primer extension. The microarray strand is attached to the SNAP array surface at the 3’ terminus. **B)** Peak overlap of the four GRHL2 ChIP-seq datasets used to construct the genomic tiling regions of the SNAP array. **C)** Close-up of a region of the full SNAP array to show varying intensities of individual bound features. GRHL2 binding is detected as red circles. **D)** Top: GRHL2 ChIP-seq peak from Reese et al. (30) with the position of the OVOL1 gene displayed. Bottom: Genomic SNAP probes tiling the ChIP-seq peak at 6-bp spacing with the normalized SNAP intensity for each probe displayed to the right. The cyan box denotes a consensus GRHL2 site on the SNAP probe. The green bases in the SNAP probes correspond to genomic sequence while the black bases correspond to the constant primer.

Representative genomic tiling across a GRHL2-bound region near the Ovo-like transcriptional repressor 1 (*OVOL1*) locus showed multiple probe variants spanning the ChIP-seq peak to identify high-and low-affinity subregions within the peak (Figure 2D). Like GRHL2, OVOL1 acts as a mediator of EMT in ovarian cancer cells (13) and exhibits a context-dependent role in breast cancer by acting as either an oncogene or tumor repressor (44, 45). Consistent with the presence of functional GRHL2 motifs, a subset of tiled probes had strong binding, whereas adjacent variants, differing only in flanking sequence or positional offset, displayed reduced fluorescence intensity. These results indicate that GRHL2 binding can be localized within ChIP-seq peaks and is sensitive to local sequence context.

Fluorescence imaging of the array confirmed discrete, reproducible binding across thousands of probe features. Bound features were distinguished from background, and the distribution of fluorescence intensities were confirmed by replicate arrays, validating the robustness of the SNAP-based affinity measurements. These data enable quantitative separation of direct, motif-driven GRHL2 binding from indirect or cofactor-mediated occupancy that is indistinguishable in ChIP-seq.

### GRHL2 binding favors the canonical AACCGGTT motif but exhibits selective tolerance to single-nucleotide variation and flanking-sequence context

The SNAP array was used to define the sequence determinants that distinguish high-affinity GRHL2 binding from partially tolerated motif variations. To do so, all genomic probe features were ranked by fluorescence intensity, and the top 1,200 highest-affinity sequences were extracted for de novo motif discovery. Across four independent bins of progressively decreasing 300 top-binding probes that total the 1,200 highest-affinity genomic sequences, MEME analysis (46) consistently recovered the canonical GRHL2 consensus motif AACCGGTT, with minimal positional variability (Figure 3A). The conservation of the consensus across the four intensity bins indicates that strong binding is dominated by recognition of the canonical octamer motif, rather than contributions from flanking or genome-context positions. To further examine the specificity of the 772,732 genomic sequences derived from the GRHL2 ChIP-seq datasets, we examined the 45,384 sequences that differed from the consensus by a single nucleotide substitution within the motif. Motif analysis of the top binders of these single mutation variants revealed a strong binding dependence on positions 1-3 and 6-8 of the motif, along with some flanking preferences when the binding window is expanded to 18 positions (Figure 3B). In a separate analysis of the flanking bases surrounding the perfect octamer motif, significant enrichment for specific 5’ and 3’ bases was observed with an optimal 5-base flanking sequence of 5’-NNNGA-AACCGGTT-CNGAG-3’ (Figure 3C). This suggests that proximal sequence environment modulates affinity without altering the core motif recognition.

**Figure 3.**
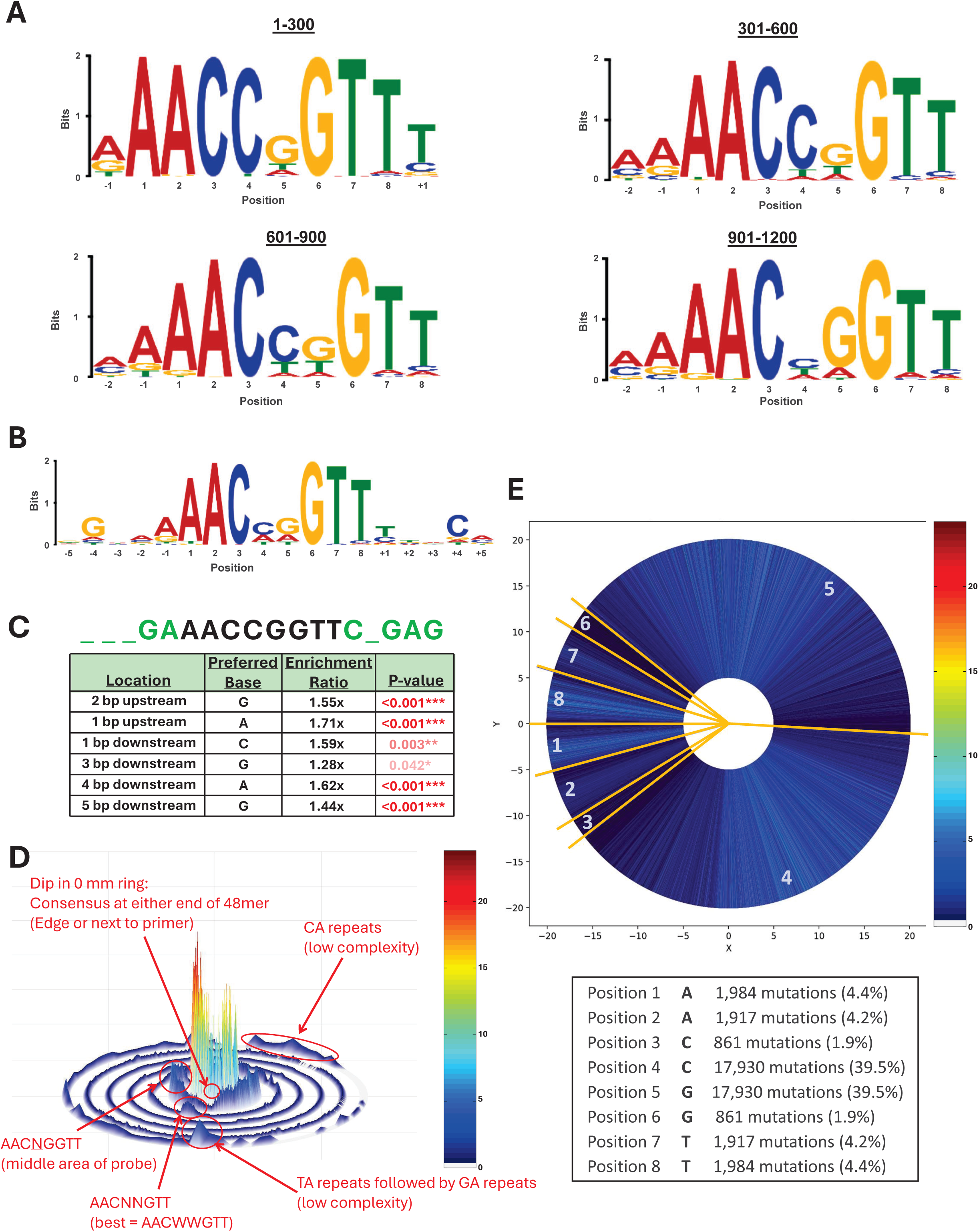
SNAP genomic binding data recovers the GRHL2 consensus motif and reveals flanking sequence preferences and mismatch distributions. **A)** Expectation maximization algorithm motifs for the highest intensity SNAP genomic sequences with a defined motif width of 10 base pairs (bp). Motif from the top 300 (top left), 301-600 (top right), 601-900 (bottom left), and 901-1200 (bottom right) highest intensity SNAP genomic sequences. Position numbering is based on the start of the GRHL2 octamer consensus, 5’-AACCGGTT-3’. **B)** Motif of the 300 highest intensity SNAP genomic sequences containing exactly one mismatch in the GRHL2 consensus motif. Five flanking bases are included in the displayed motif. **C)** Discriminatory analysis of flanking sequence surrounding the GRHL2 consensus using the highest versus lowest intensity genomic sequences. This sequence set is composed exclusively of genomic probes that contain an exact canonical GRHL2 consensus octamer, unlike **(A-B)**. Top: A summary sequence of the consensus (black) and analyzed flanking sequence (green) is provided, where underscores indicate no significant discriminatory preference. Bottom: Table listing enriched bases at specific flanking positions. The number of asterisks denote the significance of the p-value. **D)** SSL of all SNAP genomic sequences using the consensus GRHL2 motif (5’-AACCGGTT-3’) as the central SSL motif. **E)** Two-dimensional representation of the SSL 1-mismatch ring. This ring is divided into regions corresponding to where the mismatch to the GRHL2 consensus occurs. Genomically, mismatches in the central two positions of the motif occur much more commonly (∼10- to 20-fold) than the other positions. The number of probes at the differing mismatch positions is symmetrical since the motif is palindromic.

To visualize how additional sequence perturbations influence recognition, an SSL of all 772,732 sequences was constructed in which the central ring is populated with array sequences bearing the perfect AACCGGTT motif, and each successive concentric ring contains sequences that introduce one additional mismatch into the motif (Figure 3D) (34, 42). Array sequences were organized around the ring according to the specific position within the motif at which the single nucleotide substitution occurred, and then by the specific nucleotide of the mismatch. Thus, peaks in the mismatch rings correspond to specific mismatch-sequence combinations that are preferentially bound by GRHL2, providing information on both motif interdependencies as well as flanking sequence effects. Since the fluorescence intensity is proportional to the association constant (K_a_), the SSL also provides a quantitative binding energy profile relating fluorescence intensity to binding affinity (34, 42). Progressive numbers of mutations (rings 1-5) produced a graded decrease in fluorescence intensity, with the steepest decline occurring when substitutions affected the C and G at positions 3 and 6, respectively. In contrast, a large number of mismatches to the consensus sequence, which are displayed in the outer rings, produced very little binding signal. Low-complexity repeat contexts such as CA- or TA-rich flanks also modulated binding, which is consistent with some contributions from local sequence environment. This analysis revealed maximal intensity reduction by mutations at positions 3 and 6 combined with minimal intensity changes at positions 4-5 to generate a central CxxG site within the consensus AA<u>C</u>CG<u>G</u>TT motif.

To refine the tolerated variations of the motif, further analyses of the one-mismatch (Figure 3E) and two-mismatch rings were performed. The number and position of sequences around the SSL ring provide readout on base tolerances. Mutations at the C (position 3) and G (position 6) of the central CxxG were depleted among high-intensity binders as only 1.9% of the mutations, while substitutions at the terminal flanking A/T positions were more tolerated, indicating that positions 3 and 6 of the octamer contribute disproportionately to binding stability. Interestingly, mutations at the inner CG (positions 4-5) were enriched at 39.5% each, with the C most often mutated to T (C➔T: 50.2%) and A (C➔A: 46.3%), and the G conversely was most often mutated to A (G➔A: 50.2%) and T (G➔T: 46.3%). These analyses define a high-resolution positional binding landscape for GRHL2-DNA interactions, demonstrating that the canonical AACCGGTT motif is both necessary and sufficient for high-affinity binding. Furthermore, positions 3 and 6 of the octamer and shown in the central CxxG constitute the most sequence-constrained region of the site, and flanking bases modulate but do not override the octamer motif recognition.

### Extended motif analysis reveals tolerance to mismatches across dimeric GRHL2 binding sites

To determine whether GRHL2 recognizes sequence features extending beyond the canonical AACCGGTT motif, we next performed extended-motif discovery using the top 1,200 highest-intensity genomic probes of the 772,732 genomic probe set, with a constrained motif width of 18 bp. The resulting motif retained an AACCGGTT consensus but also revealed evidence of two motifs, suggestive of dimer binding on the SNAP probes (Figure 4A).

**Figure 4.**
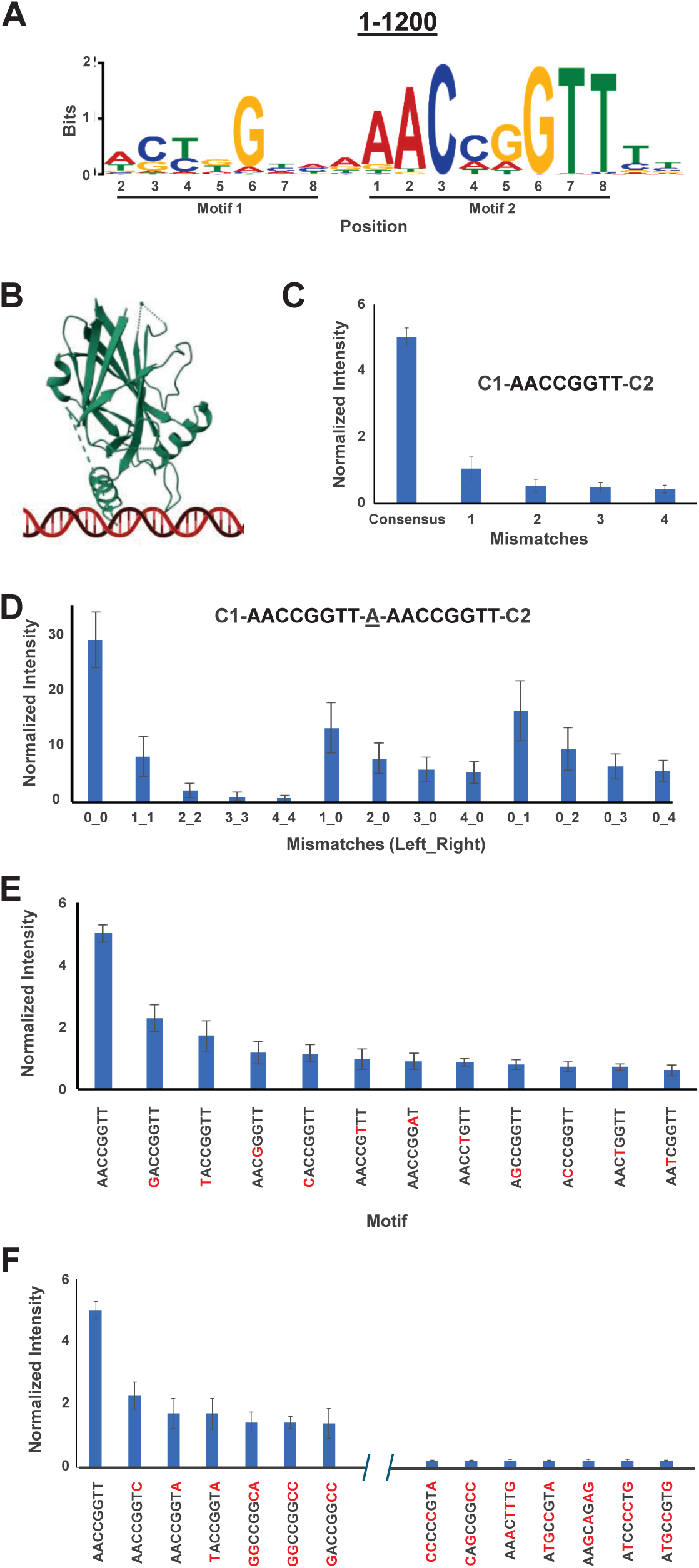
Systematic mismatch analysis reveals sequence rules governing GRHL2 monomer and dimer binding. **A)** Expectation maximization algorithm motif for the 1200 highest intensity SNAP genomic sequences with a defined motif width of 18 base pairs (bp). **B)** Crystal structure of GRHL2 DNA-binding domain (PDB: 5MR7) with overlaid linear duplex DNA schematic placed on the basis of the GRHL1-DNA crystal structure (PDB: 5MPF). **C)** Average normalized SNAP intensity for an increasing number of mismatches to the consensus sequence 5’-AACCGGTT-3’. Each mismatch group represents all possible mismatches to the consensus, resulting in 24 unique one-mismatch probes, 252 unique two-mismatch probes, 1512 unique three-mismatch probes, and 5670 unique four-mismatch probes, placed on the SNAP array in triplicate. The motifs are anchored on either side by constant regions (C1 and C2). Error bars (representing one standard deviation) are derived from all intensities of all replicate probes and possible mismatch variations within the group. **D)** Effects of mismatch(es) in a dimeric GRHL2 consensus site separated by a single base pair, flanked by constant regions. The x-axis labels denote the number of mismatches in the left binding site followed by the number of mismatch(es) in the right binding site, separated by an underscore. Each probe represented in the first five columns consists of identical mismatches in the two binding sites, while the middle and last four columns consist of mismatch(es) only in the first or second binding site, respectively, with the other binding site comprised of the consensus. **E)** Average normalized SNAP intensities for each single mismatch to the GRHL2 consensus site based on control probe intensities. Red bases denote mismatches to the GRHL2 consensus. **F)** Average normalized SNAP intensities for the six highest and seven lowest intensity mismatch motifs where the number of mismatches to the canonical GRHL2 motif vary up to four. Red bases denote mismatches to the GRHL2 consensus.

To interpret these extended preferences in the context of molecular recognition, we examined the GRHL2 DNA-binding domain crystal structure (PDB: 5MR7) overlaid with a linear duplex DNA model aligned to the GRHL1-DNA complex (PDB: 5MPF) (Figure 4B) since a crystal structure of GRHL2-DNA is not present in RSCB Protein Data Bank. This overlay is reasonable since the DNA-binding domain of these two proteins (residues 247-484 for GRHL2 from Uniprot Q6ISB3; residues 250-467 for GRHL1 from Uniprot Q9NZI5) exhibit 76% identity, 94% similarity, and a single gap of 12 residues. In this configuration, the recognition helix is positioned over the central CCGG of the octamer consensus. Residues in the wing and C-terminal extension approach the DNA phosphate backbone and adjacent flanking bases, suggesting potential protein contacts outside the core motif.

Using control probes with a motif embedded within a constant flanking sequence, we first evaluated how increasing numbers of mismatches affected binding to the consensus sequence 5’-AACCGGTT-3’ with probes bearing all possible mismatch combinations, in triplicate. Average SNAP intensities decreased as the number of mismatches increased (Figure 4C), with a sharp loss of affinity between zero and one mismatches and near-complete loss upon introduction of four mismatches. This conforms to the results of the SSL between the central and one mismatch rings (Figure 3D). These results confirm that GRHL2 binding is dependent on maintenance of the consensus sequence and that sequential disruption of the motif results in energetic penalties rather than complete loss of binding.

Based on the dimer motif (Figure 4A) and because there is evidence that the GRHL family of transcription factors can bind as dimers (43), we next examined the effects of mismatches introduced into dimeric GRHL2 motifs separated by a single base pair. Control probes were designed to contain identical or asymmetric mismatches in the left and/or right binding sites, allowing comparison of individual and multiple permutations. As shown in Figure 4D, probes containing matched mismatches in both sites displayed substantially reduced binding relative to the fully consensus dimer. However, probes with mismatches confined to a single site retained intermediate affinity. These data suggest a cooperative contribution of the paired sites to overall GRHL2 binding.

The contribution of each possible single-base substitution within the consensus site was then analyzed. Consistent with the core motif specificity described above, single mismatches at positions 3 and 6 of the central CxxG resulted in the strongest reduction in binding. Substitutions at the terminal A/T positions were more tolerated (Figure 4E and 4F). When all mismatch classes (up to four substitutions) were ranked by SNAP intensity, the variants with higher affinity maintained the central CxxG element and differed mainly at distal positions, while the lower affinity variants disrupted one or more bases within the core (Figure 4F). Interestingly, mutations in the positions flanking the central CCGG resulted in higher intensity than mutations in the middle CG (positions 4-5) as was observed in Figure 3A-B. This may be due to the fact that Figure 3 was based on genomic sequences, while the control sequences of Figure 4 evaluated all GRHL2 permutations with the identical flanking sequences. These observations reinforce that GRHL2 specificity is dominated by recognition of the central CxxG, with flanking and extended positions acting more as modulators of affinity.

### Motif spacing reveals periodic dimer-dependent binding and orientation preferences

To examine the biophysical nature of GRHL2 dimers, we further evaluated the control probes containing two GRHL2 motifs separated by increasing numbers of base pairs. We examined two configurations: a forward-forward motif (AACCGGTT-AACCGGTT) and a reverse-forward arrangement (TTGGCCAA-AACCGGTT). Fluorescence intensity varied as a function of spacing, and the pattern was distinct between the two orientations (Figure 5A). In the forward–forward configuration, binding was highest when the motifs were positioned with a 1-base spacer, with periodic decay as spacing increased. In contrast, the reverse-forward orientation preferred a 3-4 bp spacer with less apparent periodic decay.

**Figure 5.**
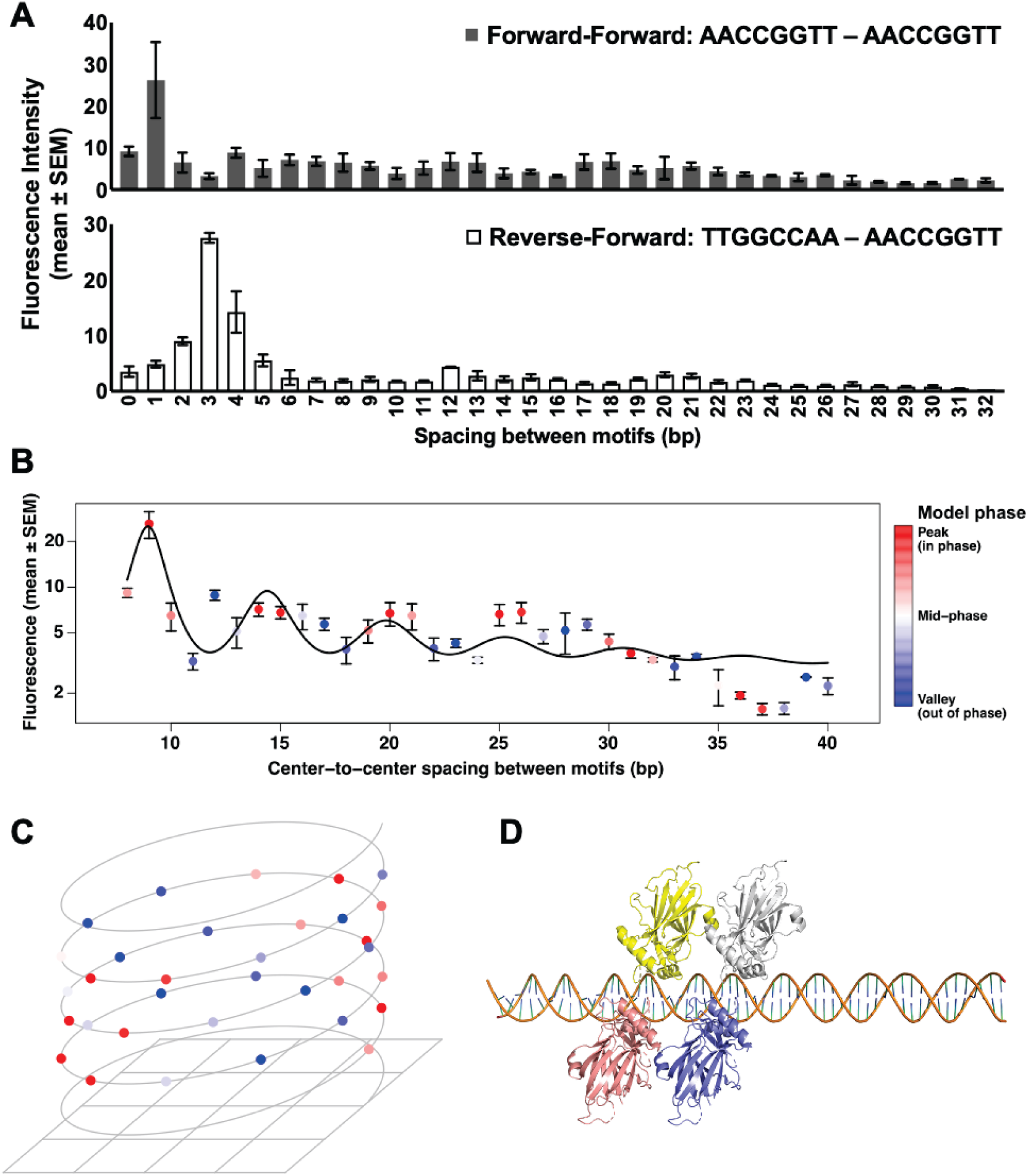
GRHL2 dimer binding strength oscillates with DNA helical periodicity. **A)** Mean fluorescence intensity (± SEM) for features containing two GRHL2 motifs separated by 0-32 bp in forward-forward (gray) or reverse-forward (white) orientation, illustrating orientation-dependent differences in binding signal. **B)** Forward-forward fluorescence values plotted on a logarithmic scale as a function of center-to-center motif spacing. Points represent mean ± SEM. The black curve shows the best-fit torsional phasing model, yielding periodicity of approximately 5.46 bp. Data points are colored by the phase of the fitted helical periodicity model. **C)** Helical representation of the forward-forward data, mapping motif spacing onto the expected geometry of the DNA helix (10.5 bp per turn). Points retain color coding from panel **(B)**, highlighting that peak binding intensities cluster on alternating faces of the DNA helix, consistent with preferential binding when GRHL2 dimers occupy the same helical face. **D)** Structural model illustrating the predicted configuration at the optimal forward-forward spacing of 1 bp. Using the crystal structure of GRHL1 dimer bound to DNA, two GRHL2 dimers, depicted as yellow and salmon monomers forming dimer 1 and white and blue monomers forming dimer 2, were positioned to model binding to two DNA motifs spaced 1 bp apart.

Because the higher intensity was observed in the forward-forward motif, we analyzed this configuration in more detail (Figure 5B). Plotting fluorescence intensity on a logarithmic scale against spacing revealed a repeating pattern, with peaks occurring at regular intervals. Curve fitting yielded a periodicity of approximately 5.5 bp, corresponding to approximately half of the 10.5-bp helical repeat of B-form DNA (38). Data points were color-coded according to their phase within the fitted helical phasing model, and this showed that higher binding tends to occur when the two motifs align on the same rotational face of the helix.

To visualize this effect in terms of DNA geometry, the spacing arrangement was projected onto a helical representation (Figure 5C). The colored peak positions clustered on alternating faces of the helix, indicating that GRHL2 dimers bind most strongly when both sites fall on the same helical surface rather than being rotated away from one another. This pattern supports a model in which spatial positioning along the helix facilitates simultaneous or cooperative binding of two GRHL2 molecules.

A structural model was generated to illustrate the GRHL2 configuration predicted at the optimal spacing distance (Figure 5D, Supplementary Movie 1). Using the published crystal structure of a GRHL1 single dimer (comprised of two monomers) on a single DNA motif as a template (43), two GRHL2 dimers (each comprised of two monomers) were positioned on paired motifs separated by each spacing of increasing base pairs using PyMol. The intervening B-DNA fiber structures used to define precise helical geometry and spacing were generated with Web 3DNA 2.0 (47). At 1-bp spacing, the two sets of GRHL2 dimers are oriented on the same side of the DNA in close proximity (Figure 5D). These results suggest that GRHL2 can bind dimeric AACCGGTT sites with preference for those that are arranged in the same orientation and positioned on the same helical face, which may reflect geometric constraints of dimer-dimer interactions on DNA.

### Distinguishing direct GRHL2 binding from indirect, cofactor-mediated recruitment across genomic loci

The GRHL2 genomic SNAP array was next used to discriminate genomic regions where GRHL2 binds DNA directly from those where ChIP-seq enrichment likely reflects indirect, protein-tethered interactions (Figure 6A). If a genomic region showed GRHL2 occupancy by ChIP-seq and also produced a detectable signal on the SNAP array, the site was classified as a direct DNA-binding event. In contrast, regions with strong ChIP-seq signal but no measurable binding on the SNAP array were considered indirect binding events, consistent with GRHL2 being tethered through an interacting transcription factor or cofactor rather than contacting DNA directly at the motif. Representative examples of each category of binding event are shown in Figure 6B. At the *MCTS1* promoter, GRHL2 ChIP-seq peaks were observed under both endogenous and overexpressed GRHL2 conditions (7, 30), and the SNAP array revealed a clear binding signal across a subset of adjacent tiled probes in this region that represent a single GRHL2 binding site approximately at the center of the ChIP-seq peak, indicating direct recognition of underlying DNA sequences. At the *EPIC1* promoter, GRHL2 was also enriched by ChIP-seq. However, SNAP probes spanning the same interval showed little to no binding signal, consistent with GRHL2 localization being mediated by another DNA-bound factor at this site.

**Figure 6.**
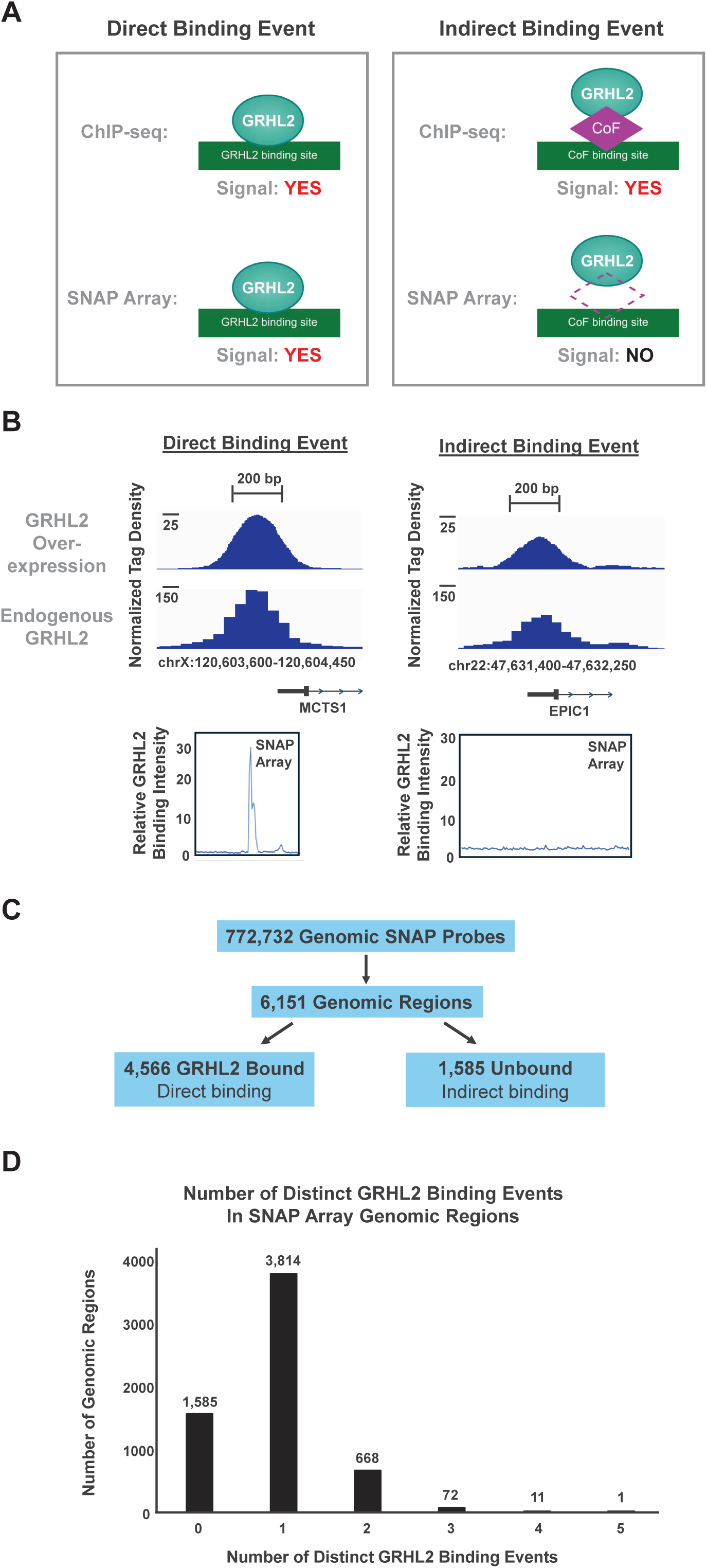
SNAP data distinguished direct GRHL2 DNA binding from indirect protein-mediated recruitment at genomic sites. **A)** Schematic demonstrating the method to determine direct versus indirect binding of GRHL2 at genomic sites. Left: When binding is observed by both ChIP-seq and SNAP arrays at the same genomic location, this is considered a direct binding event. Right: When binding is observed by ChIP-seq but not SNAP arrays, then the ChIP-seq binding is considered to be mediated by a secondary cofactor (CoF) that indirectly tethers GRHL2 to this genomic location. **B)** Representative examples of a direct (Left) and indirect (Right) binding event. ChIP-seq signal from overexpressed (Top) and endogenous (Middle) GRHL2 levels at the MCTS1 and EPIC1 promoters (7, 30). Normalized GRHL2 binding intensity (Bottom) is shown at these promoters as tiled on the SNAP array. Each region contains multiple DNA probes tiled across the region. **C)** Summary of the genomic probe count on the SNAP array with the corresponding number of genomic regions evaluated using overlapping SNAP probes. These genomic regions are then divided into direct and indirect GRHL2 sites based on the SNAP probe in each region with the highest intensity. **D)** Bar graph of the number of distinct GRHL2 binding sites within each genomic region tiled by the SNAP array. One region, chr7:155,123,400-155,125,400 (Figure 7), was excluded from this analysis due a large number of direct binding sites within a very short region, where numerous SNAP probes contain multiple GRHL2 direct binding sites.

Across the full genomic SNAP dataset, 772,732 probes collapsed into 6,151 distinct genomic regions based on tiling overlap (Figure 6C). Using the highest-intensity probe within each region as the representative value, 4,566 regions were classified as direct GRHL2 binding sites (which includes perfect and mismatch sites), while 1,585 regions showed ChIP-seq signal without SNAP binding and were therefore assigned as indirect sites. These data indicate that 25.8% of GRHL2 ChIP-seq peaks tiled on the SNAP array correspond to indirect recruitment events rather than direct recognition of the DNA motif. As ERa and FoxA1 are transcription factors reported to have direct interactions with GRHL2, the 1,585 indirect binding regions were compared to the 4,566 direct binding regions in terms of both the genomic sequence and ChIP-seq datasets for these transcription factors (29, 48, 49). ERa did not have any statistical difference in the relative number of regions that contain a genomic ERa half-site (5’-RGGTCA-3’ half-site where R = A or G) nor in the relative number of regions that contain a ChIP-seq peak from two different studies of ERa in the MCF7 breast cancer cell line (Supplementary Table 1). The ERa ChIP-seq data were conducted with antibodies against total ERa protein (49) or specifically against phosphorylated ERa at S118 (29). In contrast, the indirect regions exhibited a higher relative proportion of FoxA1 ChIP-seq binding sites as compared to direct regions, suggesting that recruitment of GRHL2 to the indirect regions may be driven, in part, by protein tethering with FoxA1 rather than ERa.

The number of distinct direct binding events occurring within each genomic interval tiled on the array was then determined. Most regions contained either one or two distinct direct GRHL2 binding sites, with a smaller number containing three or more (Figure 6D). Overall, these results demonstrate that combining SNAP array measurements with genome-wide ChIP-seq studies enables functional separation of direct sequence-encoded GRHL2 binding from cofactor-dependent occupancy, revealing distinct classes of GRHL2-associated genomic loci.

### A dense cluster of GRHL2 motifs corresponds to a highly occupied ChIP-seq region

In examining how often the GRHL2 binding motif occurs across the genome, a low-complexity region on chromosome 7 that contains an unusually high density of GRHL2 consensus motifs was identified (Figure 7A). Within this interval from chr7:155,124,004-155,124,759, multiple exact canonical AACCGGTT motifs were present, interspersed with additional single-mismatch variants. This arrangement created a concentrated cluster of closely spaced GRHL2 recognition sites across a relatively short stretch of DNA. ChIP-seq data from our group (7, 30) showed strong GRHL2 enrichment at the edges of this region under both datasets (Figure 7B, top). When mapped on the SNAP array, probes tiled every six base pairs across the same interval produced a broad band of elevated binding signal within the central region (Figure 7B, bottom). Rather than a single discrete binding peak, the SNAP signal fluctuated across the tiled probes, which is consistent with multiple binding events arising from the repeated consensus and near-consensus motifs. The observed difference between the ChIP-seq and SNAP data could be explained by imprecise mapping of ChIP-seq reads in this low-complexity region that resulted in two peaks on either side of the region. In particular, the ChIP-seq peak on one side of the region may represent the perfect GRHL2 consensus (5’-AACCGGTT-3’) reads while the ChIP-seq peak on other side of the region may represent the primary mismatch site (5’-AAACGGTT-3’) reads. Alternatively, this region may be chromatin inaccessible to the ChIP-seq antibody reagents. These observations indicate that this region potentially represents a highly occupied GRHL2 locus driven by the presence of many closely spaced binding sites. The combination of dense consensus motifs and favorable single-mismatch variants likely creates a local DNA environment that strongly favors GRHL2 binding across the entire interval.

**Figure 7.**
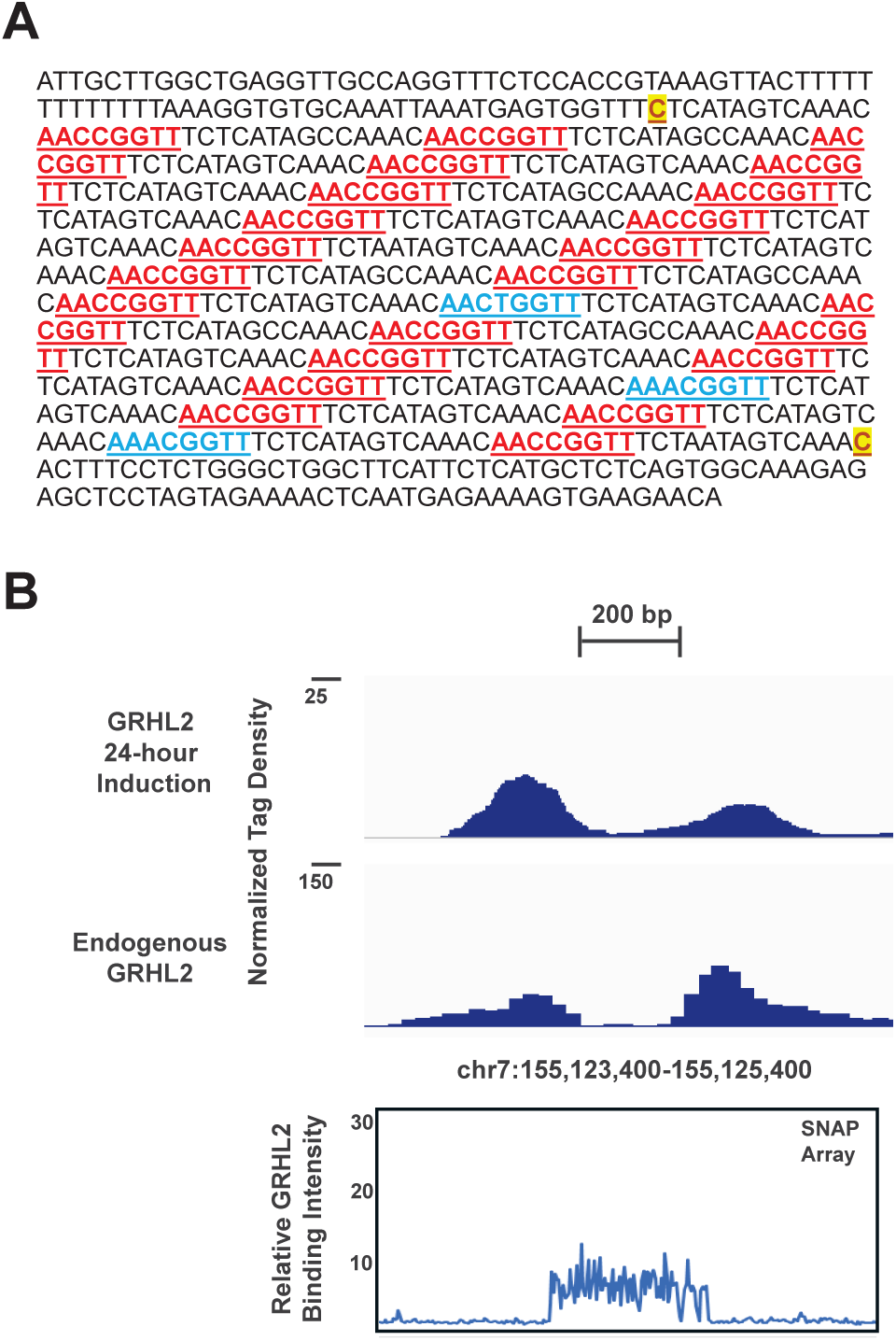
A genomic region with extreme GRHL2 motif density produces broad SNAP binding signal that is not fully reflected in ChIP-seq profiles. **A)** Genomic sequence of chr7:155,124,004-155,124,759 (GRCh38). Consensus motifs are underlined with perfect sites in red and single mismatch sites in blue. Low complexity repeats of 22 bp start and end with the bases in yellow highlight. **B)** Top: ChIP-seq data from Zheng et al. (7) and Reese et al. (30) are displayed for this region. Bottom: Normalized SNAP intensities for genomic probes across this region tiled at every 6 bp are displayed for this region.

## DISCUSSION

In this study, we used a customized high-density genomic SNAP DNA-binding array to biochemically define, at six-base resolution, the sequence features that govern GRHL2 DNA recognition across the genome. By integrating direct binding measurements with structural interpretation and ChIP-seq data, we distinguish motif-encoded GRHL2 direct binding events from cofactor-dependent occupancy to resolve how motif composition, flanking sequence, and motif spacing collectively dictate GRHL2 binding characteristics. While prior studies have established GRHL2 as a co-regulator of transcription through interactions with factors such as ERa and PR and through modulation of chromatin accessibility, these approaches do not differentiate direct DNA binding from indirect, cofactor-mediated recruitment. The present work addresses this gap by defining the intrinsic sequence determinants of GRHL2 binding independent of chromatin context (50, 51).

Across more than 772,732 genomic probe sequences, GRHL2 binding prefers a conserved AACCGGTT motif. Substitutions at positions 3 (C) and position 6 (G) produced the steepest reductions in SNAP intensity. Substitutions at the central CG (positions 4-5) had minor effects, and substitutions at the outer A/T positions were more tolerated. These results are consistent with structural studies of the Grainyhead/CP2 family (43). The recognition helix of the winged-helix DNA-binding domain contacts the positions 3 and 6 of the AACCGGTT motif with no direct contacts to the central CG, resulting in a CxxG core. The limited backbone contacts outside of the octamer core motif found in the crystal structure (43) suggest that the flanking preferences uncovered in our analysis may be driven by DNA structural features, such as major groove geometry influenced by A/T content. The SSL and mismatch analyses also exhibited this enrichment of tolerated single-base substitutions at the middle CG positions of the motif, with a strong bias toward C➔T/A and G➔A/T changes. One possible explanation is that these substitutions preserve DNA shape or hydrogen-bonding geometry in a way that is partially compatible with GRHL2 recognition, allowing low-affinity binding rather than the complete loss of the interaction.

The SSL analysis provides quantitative detail towards a structural model in which the central CxxG acts as the principal energetic contributor for GRHL2-DNA recognition, while flanking sequences and the central CG tune binding affinity and specificity. This level of resolution extends beyond traditional motif enrichment analyses of ChIP-seq peaks by providing a quantitative description of exactly where GRHL2 may bind within a genomic region to a “sub-peak” level. This enables direct interpretation of genomic GRHL2 occupancy to distinguish sequence-driven binding from indirect recruitment observed in chromatin-based assays.

The enrichment of tolerated substitutions at the central CG positions further suggests that GRHL2 binding may be relatively robust to sequence variation at CpG dinucleotides, which are known to be highly mutable in mammalian genomes due to cytosine deamination (13). The observed tolerance to specific C➔T/A and G➔A/T substitutions brings forth the idea that GRHL2 binding can persist, at reduced affinity, in regulatory regions undergoing CpG erosion. This may have implications for the evolution of GRHL2 binding sites and for the maintenance of regulatory function in genomic regions that are known to be mutationally active.

Our spacing and orientation analyses demonstrate that GRHL2 binding is affected by motif organization. Paired motifs in a forward–forward orientation (AACCGGTT-AACCGGTT) exhibited the strongest binding at short distances and displayed a periodic pattern with an approximate 5.5-bp interval, consistent with half of the DNA helical turn (38). This periodicity suggests that optimal binding occurs when motifs are positioned on the same face of the DNA helix, which supports a model of cooperative or dimer binding. While previous studies have suggested that members of the Grainyhead family can form dimers, the present results define the spatial constraints of these interactions and provide a structural basis for motif clustering observed within GRHL2-bound regulatory regions. The dense motif cluster on chromosome 7, where repeated consensus and near-consensus motifs produced a broad region of elevated SNAP signal and ChIP-seq occupancy, provides a natural genomic example of how such multivalent binding environments may occur in the genome.

These findings provide a method for interpreting GRHL2 co-occupancy with other transcription factors in genomic studies. Recent work has demonstrated that GRHL2 co-binds enhancer regions with PR and contributes to chromatin accessibility and transcriptional regulation in hormone-responsive breast cancer cells (50). However, such approaches cannot resolve whether GRHL2 is directly engaging DNA at these sites or is recruited through protein-protein interactions. By distinguishing direct motif-driven binding from indirect occupancy, our results act in complement with genome-wide studies to show that only a subset of GRHL2 ChIP-seq peaks represent sequence-encoded binding events, while others may reflect cofactor-mediated recruitment. This distinction has important implications for interpreting genomic occupancy data and for identifying functionally relevant GRHL2 binding sites. This observation is consistent with prior transcriptomic and cistromic studies showing that GRHL2 interacts with ERa, FoxA1, and other transcriptional regulators and frequently participates in enhancer environments rather than acting as a solo DNA-binding factor (29, 30, 50, 51).

Towards potential implications of these findings for breast cancer biology, GRHL2 has been identified in maintaining epithelial identity and in modulating ERa-dependent transcriptional programs, yet it has also been associated with worse clinical outcomes in ER-positive tumors. The fine-scale binding specificity patterns described here suggest that GRHL2 occupancy may be highly sensitive to sequence perturbations within the central CxxG or to cooperative spacing relationships at dimeric sites. Somatic mutations affecting these positions or CpG instability at GRHL2-enriched regions could shift GRHL2 binding patterns in tumors, potentially weakening epithelial maintenance programs or altering ER-associated enhancer activity. Conversely, regions that retain strong GRHL2 binding capacity may reinforce epithelial transcriptional states and could represent distinct therapeutic response contexts. Although our study does not directly assess these relationships, the binding parameters established here provide information towards mapping how patient-specific sequence variation may remodel the GRHL2 cistrome.

By analyzing GRHL2 binding at ChIP-seq sites in the absence of chromatin and cooperative transcriptional co-regulators, this work provides a compendium of GRHL2 binding data for 770,000-plus genomic sites and defines, at an unprecedented high resolution, the sequence specificity underlying GRHL2 DNA recognition across genomic sequence space. The ability to resolve binding at sub-peak resolution, quantify the effects of sequence variation, and define spacing-dependent interactions advances our understanding of GRHL2 beyond consensus motif models toward a predictive, mechanistic description of its DNA-binding specificity. Our findings show that GRHL2 binding is anchored by a highly constrained central CxxG within its octamer consensus motif and is affected by nearby flanking sequence outside of its consensus. Our results support that GRHL2 binding is strengthened at dimeric sites and can be distinguished as either direct DNA binding or cofactor-mediated occupancy across the genome. These insights refine the structural and biochemical understanding of GRHL2 interaction with DNA and lay the groundwork for future studies linking CpG mutational landscapes with GRHL2-dependent transcription in development and disease, as well as methods to disrupt key GRHL2 binding sites using site-directed therapeutics.

## ACKNOWLEDGEMENTS

The authors thank members of the Fowler Laboratory, Alarid Laboratory, and Proteovista LLC for their helpful comments on the data and manuscript. In particular, we thank Natasha Solodin, Blue-leaf Cordes, Kelley Salem, Kaelyn Allen, and Christy Zheng for scientific discussions.

## AUTHOR CONTRIBUTIONS

Conceptualization: Mary S. Ozers, Elaine T. Alarid, Christopher L. Warren, Paige E. Messa, Keenan S. Pearson

Data Curation: Paige E. Messa, Christopher L. Warren, Noah R. Nicol

Formal Analysis: Christopher L. Warren, Paige E. Messa, Noah R. Nicol, Keenan S. Pearson, Justin P. Peters

Funding acquisition: Elaine T. Alarid, Amy M. Fowler, Mary S. Ozers

Investigation: Paige E. Messa, Keenan S. Pearson, Christopher L. Warren, Noah R. Nicol Methodology: Paige E. Messa, Keenan S. Pearson, Christopher L. Warren, Noah R. Nicol, Mary S. Ozers

Project Administration: Mary S. Ozers, Elaine T. Alarid Resources: Mary S. Ozers

Software: Noah R. Nicol, Christopher L. Warren Supervision: Mary S. Ozers, Elaine T. Alarid, Amy M. Fowler

Validation: Paige E. Messa, Keenan S. Pearson, Christopher L. Warren, Noah R. Nicol Visualization: Paige E. Messa, Keenan S. Pearson, Christopher L. Warren, Noah R. Nicol, Mary S. Ozers

Writing – original draft: Paige E. Messa, Mary S. Ozers

Writing – review & editing: Paige E. Messa, Christopher L. Warren, Noah R. Nicol, Keenan S.

Pearson, Justin P. Peters, Amy M. Fowler, Elaine T. Alarid, Mary S. Ozers

