## Supplementary Data for "Defining the DNA Binding Specificity of GRHL2"

**Supplementary Figure 1**

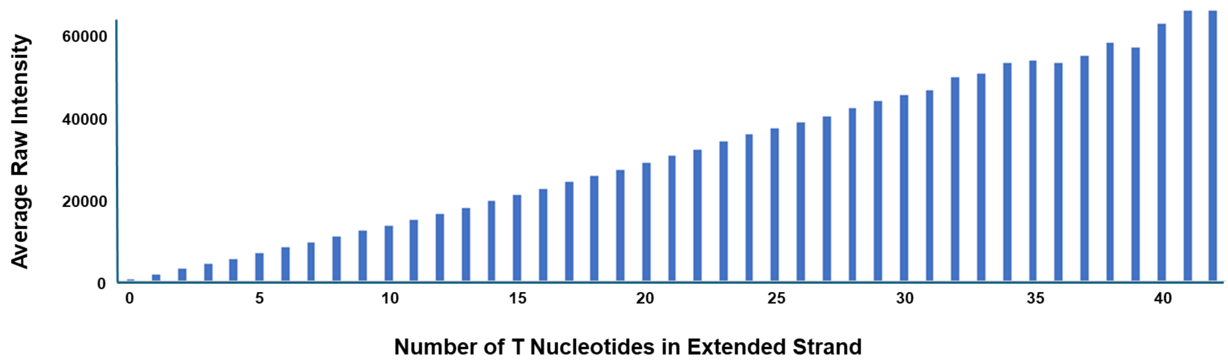


**Supplementary Figure 1. Primer extension efficiency on the SNAP array was validated by proportional Cy3-dUTP signal**. In a representative array, a linearly increasing intensity is observed with a corresponding increase in the number of T nucleotides incorporated in the extended strand (excluding the primer sequence), where the raw fluorescent signal is obtained from Cy3-dUTP.

**Supplementary Movie 1**

**A)**

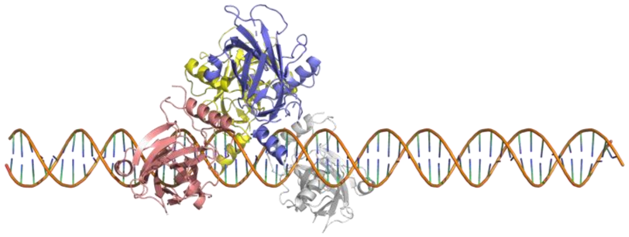

**B)**

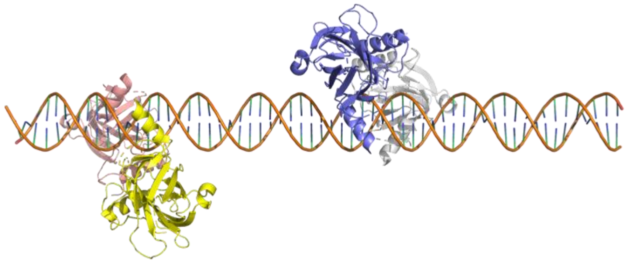

**C)**

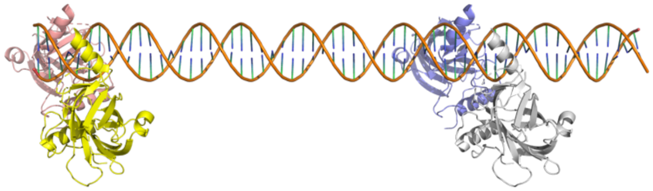

**Supplementary Movie 1.** Structural model showing predicted configurations of two GRHL2 dimers across all tested DNA motif spacings between two forward-forward motifs (0-32 bp). The 1 bp forward–forward spacing, which yielded the highest fluorescence and is shown in Figure 5D, is included among the modeled conformations. Still images are shown as A) beginning, B) middle, and C) end of the Supplementary Movie 1 file at bp spacings of A) 0, B) 16, and C) 32.

**Supplementary Table 1**

| ChIP-seq | | | | | | Unique SNAP Regions | | Chi-Squared |
|---|---|---|---|---|---|---|---|---|
| GEO | Reference | Author | Protein | Antibody | Total Peaks | Direct Overlap | Indirect Overlap | p-value |
| GSE112969 | PMC6408623 | Carroll JS | FoxA1 | ab-23738 | 119,080 | 975 | 484 | $1.32 \times 10^{-8}$ |
| GSE128208 | PMC6457525 | Siersbaek R | ERa_total | sc-543 | 6,046 | 23 | 14 | 0.138 |
| GSE117569 | PMC6336141 | Alarid ET | ERa_total | sc-543 | 56,224 | 132 | 48 | 0.853 |
| GSE117569 | PMC6336141 | Alarid ET | ERa_pS118 | ab-32396 (ab2) | 5,041 | 12 | 5 | 0.948 |
| GSE117569 | PMC6336141 | Alarid ET | ERa_pS118 | sc-12915 (ab3) | 23,052 | 60 | 30 | 0.132 |

**Supplementary Table 1. GRHL2 indirect SNAP binding sites associate near known genomic FoxA1 binding sites.** A statistical difference in the proportion of total ChIP-seq peaks from datasets that overlapped with direct versus indirect SNAP regions was evaluated using the chi-squared test of independence. ChIP-seq assays in GSE117569 were performed in triplicate, and therefore average counts were used in each column for these experiments. Only FoxA1 (GSE112969) exhibited a statistically significant difference ($p < 0.05$). Since there are an overall total of 4,566 direct and 1,585 indirect SNAP regions, this result indicates that there are a larger than expected number of FoxA1 ChIP-seq peaks overlapping with the indirect SNAP regions relative to the number of FoxA1 ChIP-seq peaks that overlap with direct SNAP regions.
